# Intestinal-epithelial LSD1 controls cytoskeletal-mediated cell identity including goblet cell effector responses required for gut inflammatory and infectious diseases

**DOI:** 10.1101/2020.07.09.186114

**Authors:** Naveen Parmar, Kyle Burrows, Håvard T. Lindholm, Rosalie T. Zwiggelaar, Mara Martín-Alonso, Madeleine Fosslie, Bruce Vallance, John Arne Dahl, Colby Zaph, Menno J. Oudhoff

## Abstract

Infectious and inflammatory diseases in the intestine remain a serious threat for patients world-wide. Reprogramming of the intestinal epithelium towards a protective effector state is important to manage inflammation and immunity. The role of epigenetic regulatory enzymes within these processes is not yet defined. Here, we use a mouse model that has an intestinal-epithelial specific deletion of the histone demethylase *Lsd1* (cKO mice), which maintains the epithelium in a fixed reparative state. Challenge of cKO mice with chemical-induced colitis, bacteria-induced colitis, and a helminth infection model all resulted in increased pathogenesis. Mechanistically, we discovered that LSD1 directly controls genes that facilitate cytoskeletal organization, and that this is relevant for epithelial attachment as well as for goblet-cell specific effector responses.This study therefore identifies intestinal-epithelial epigenetic regulation by LSD1 as a critical element in host protection from inflammation and infection.

## INTRODUCTION

Gastrointestinal infections remain one of the most common causes for hospitalization worldwide (Kirk et al., 2015). These infections can be caused by diverse pathogens including bacteria, viruses, and parasites. Despite recent advances in treatment for infections, the increase in incidence of antibiotic resistant bacterial strains likely will ensure that infection remains a large threat to global health (Carlet, 2012). Therefore, increasing our understanding of disease mechanisms and exploring new options such as host-directed therapeutics remains a very important focus in basic research.

The intestinal epithelium is a crucial component of the gut barrier and is composed of a single layer of intestinal epithelial cells (IECs) including enterocytes, Paneth cells, goblet cells and tuft cells (Allaire et al., 2018). Several of these cell types are involved in the protection against pathogens. For example, Paneth cells that reside at the bottom of small intestinal crypts are a source of antimicrobial proteins such as lysozyme and α-defensins (Bevins and Salzman, 2011). Likewise, goblet cells secrete mucins and antimicrobials that are effective against bacteria and protozoa (Kim and Ho, 2010). On the other hand, tuft cells express IL-25 that is important in mounting a type 2 immune response to protect against certain helminths (Ting and von Moltke, 2019). Finally, different pathogens elicit different immune responses that result in an alteration of the intestinal epithelial cell composition that is important for effector responses (Biton et al., 2018; Lindholm et al., 2020). These epithelial effector responses are ultimately critical for immunity against intestinal pathogens.

In addition to epithelial effector mechanisms targeting pathogens, there is also a damage response upon inflammation or infection. There is a sequence of events that include migration and proliferation of epithelial cells, which eventually results in the re-establishment of mucosal homeostasis (Benz et al., 2017). To regain epithelial integrity after an insult, IECs are reprogrammed into a temporal reparative state that is required for repair (Wang et al., 2019; Yui et al., 2018). Mice that lack the ability to reprogram towards such reparative state succumb from an otherwise manageable colitic insult (Cai et al., 2010; Yui et al., 2018). However, it is unknown whether a reparative epithelial state prior to inflammation or infection is beneficial for the host.

Chromatin accessibility has an important role in regulating gene expression. Chromatin accessibility can be mediated by posttranslational modifications of DNA and Histone proteins, which is done by epigenetic modifiers (Kouzarides, 2007). In the intestinal epithelium, the importance of maintaining epigenetic marks is exemplified by the finding that loss of the polycomb repressor complex 2 (PRC2) leads to a rapid loss of various cell types including intestinal stem cells, which subsequently leads to loss of tissue integrity and mice become moribund (Chiacchiera et al., 2016; Jadhav, 2017; Koppens et al., 2016). We recently found an important role for the demethylase LSD1 in early-life intestinal epithelial development, including the differentiation of Paneth cells (Zwiggelaar et al., 2020). LSD1 can demethylate mono or di-methylated Histone 3 Lysine 4 (H3K4me1/2) that normally mark active or poised enhancers (Shi et al., 2004). Importantly, we previously found that LSD1-deficient small intestinal epithelium is in a continuous state of repair (Zwiggelaar et al., 2020).

## RESULTS

### LSD1-deficient epithelium is in a reparative state during homeostasis

We have recently shown that inhibition or deletion of the histone demethylase LSD1 renders the small intestinal epithelium in a continuous state of repair, and this state is beneficial for recovery after irradiation injury (Zwiggelaar et al, 2020). We generated mice that lack *Lsd1* in the intestinal epithelium specifically by crossing *Villin-*Cre mice with *Lsd1*^f/f^ mice (conditional knock-out (cKO) mice) and found that these mice have a near complete deletion of *Lsd1* in small intestinal epithelium (Zwiggelaar et al., 2020). In the current study, we focus on the colon, and here we also find crypts still expressing LSD1 in cKO mice (Fig. S1A). Nevertheless, we assessed whether the colonic epithelium in the cKO mice is in a general reparative state by examining different parameters such as proliferation and presence of the reparative marker Sca-1. Although we do not observe a significant increase in crypt length, we did find increased proliferation in colonic crypts of cKO mice compared to *Lsd1*^f/f^ (WT) littermates as measured by Ki67^+^ cell counts (Fig. 1A, 1B). In support, in organoids, we found that the Ki67^+^ proliferative zone was expanded and with increased intensity in cKO organoids compared to WT organoids (Fig. 1C, 1D). Thus, increased proliferation is an intrinsic effect of *Lsd1* deletion in intestinal epithelium. Two studies identified Sca-1 as a bona fide marker of repairing epithelium, and Sca-1-derived organoids partially retain this reparative state (Nusse et al., 2018; Yui et al., 2018). To test whether *Lsd1*-deficient epithelium contains Sca-1^+^ cells, we isolated colon crypts and analyzed Sca-1^+^ cells by flow cytometry. Indeed, we observed that crypt cells derived from cKO mice contain more Sca-1^+^ cells than their WT cage littermates (Fig. 1E, 1F, S1B). Finally, IL-22 is known to be a crucial cytokine in the defense against pathogens, which among other responses is due to inducing epithelial repair (Lindemans et al., 2010), and by inducing antimicrobials such as REG3B and REG3γ (Zheng et al., 2008). To test if inhibition of LSD1 would affect the IL-22-induced antimicrobial responses, we co-treated WT organoids with a potent LSD1 inhibitor that phenocopies *Lsd1* deletion (Zwiggelaar et al., 2020) and IL-22. We found that IL-22 induced STAT3 Tyr705 phosphorylation by western blot and confocal imaging, and robustly induced *Reg3b* and *Reg3g* by qPCR independently of LSD1 enzymatic activity (Fig. 1G, 1H, S1C). Together, these findings suggest that the colon epithelium of cKO mice is in a continuous reparative state and that this reparative state does not hamper IL-22-induced antimicrobial programs.

**Fig. 1.**
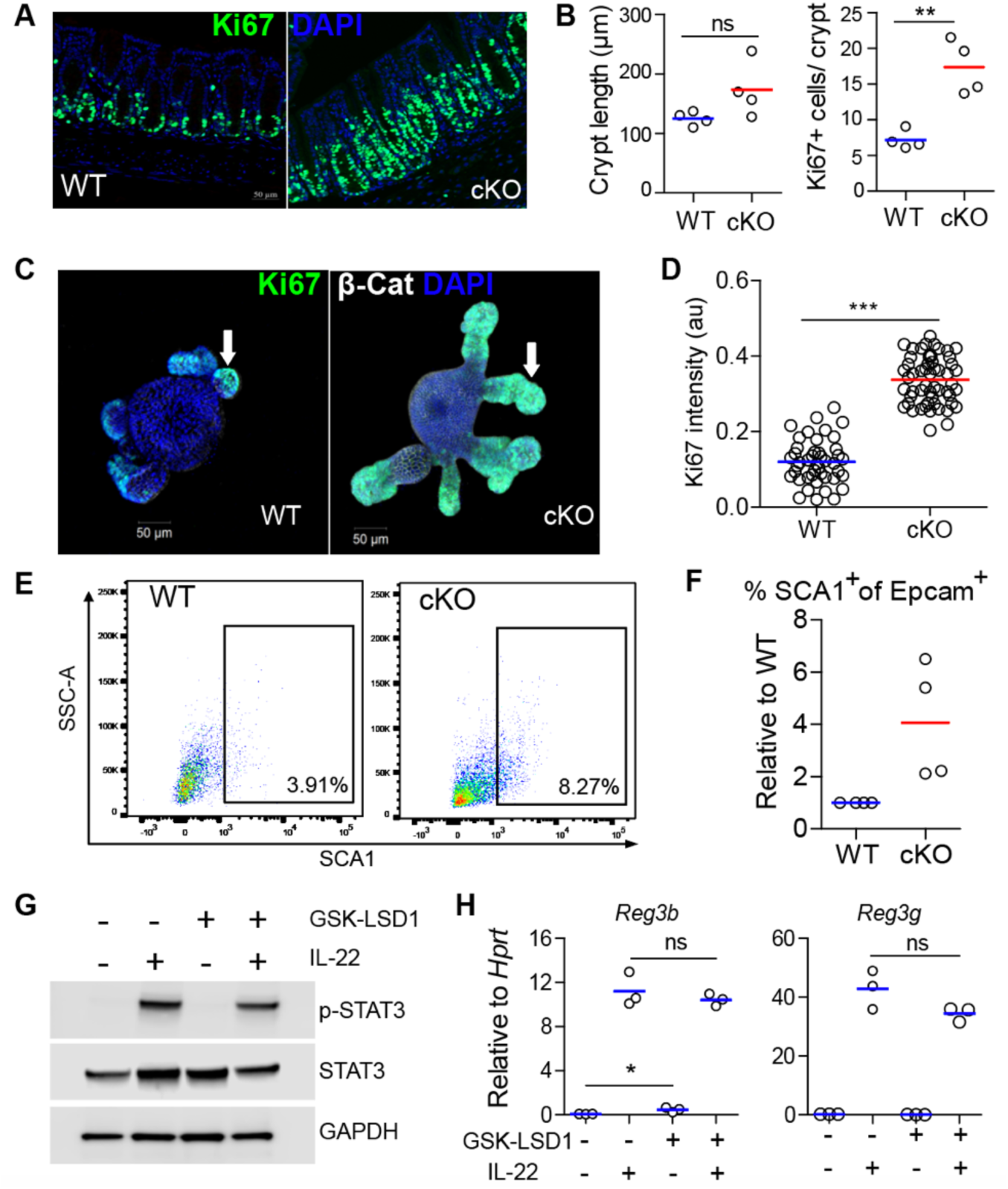
Intestinal epithelial *Lsd1* deletion induces proliferation and a repair-like state in epithelium of mice. **A**, Immunofluorescent images of Ki67**+** (green) in colonic epithelium of WT and cKO adult mice. DAPI was used as a nuclear counterstain. **B**, Quantification of crypt length in naïve WT and cKO adult mice. Quantification of Ki67**+** cells in colonic crypts of WT and cKO adult mice. n= 4 mice in each group, line is mean. **C**, Organoids from WT and cKO mice were cultured for 3 days. The expression of Ki67+ (green) was determined by immunofluorescence. **D**, Quantification of Ki67 intensity in budding organoids from WT and cKO mice. Each dot represents 1 organoid. **E**, Crypts cells were isolated from colon of WT and cKO adult mice and flow cytometric analysis of SCA1 was carried out. **F**, Quantification of SCA1**+** positive cells in crypts from WT and cKO adult mice, lines indicate littermate controls. n= 4 mice **G**, Organoids were treated with GSK-LSD1 for 4 days followed by IL-22 stimulation for 30 minutes. Whole cell lysates were prepared and subjected to immunoblot analysis of p-STAT3, STAT3, and GAPDH. **H**, Organoids were co-cultured with GSK-LSD1 (5μM) and IL-22 (10 ng/ml) for 4 days in the ENR medium. The mRNA expression of *Reg3b* and *Reg3g* was determined by RT-qPCR relative to housekeeping gene *Hprt*. n = 3 biological replicates of a representative experiment. Unpaired two-tailed Student’s t test (B,D & H) was performed to observe significant differences. ns = not significant, * P ≤ 0.05, ** P < 0.01, *** P ≤ 0.001.

### LSD1 is protective during chemically induced colitis

Inflammatory bowel disease represents two major types of intestinal disorders that causes chronic inflammation of the digestive tract (Rubin et al., 2012). In our study, we assess the severity of colonic inflammation after administration of dextran sulfate sodium (DSS, 3.5 %) comparing WT and cKO cage littermates. Yui et al. identified that YAP/TAZ-mediated reprogramming of intestinal epithelium into a repair state is important in this disease model (Yui et al., 2018), and we found that *Lsd1*-deficient epithelium matches the phenotype of reparative epithelium (Fig. 1, and (Zwiggelaar et al., 2020). We thus wanted to test whether *Lsd1*-deficient epithelium would be protective upon inflammatory damage, like it is upon damage induced by irradiation. In stark contrast to our previous findings using irradiation, we observed that DSS in drinking water resulted in aggravation of colonic inflammation as indicated by increased mortality (males) and decrease in body weight (both males and females) in cKO mice as compared to their WT littermates (Fig. 2A, 2B). In support, cKO mice had shorter colons compared to WT mice (Fig. 2C). Moreover, histological analysis demonstrates that damage and inflammation was more progressive in cKO mice in comparison with WT mice (Fig. 2D, 2E). In addition, we observed regions with withered crypts in cKO tissue (Fig. S2A). Therefore, we conclude that intestinal-epithelial LSD1 has a protective role during acute intestinal inflammatory responses.

**Fig. 2.**
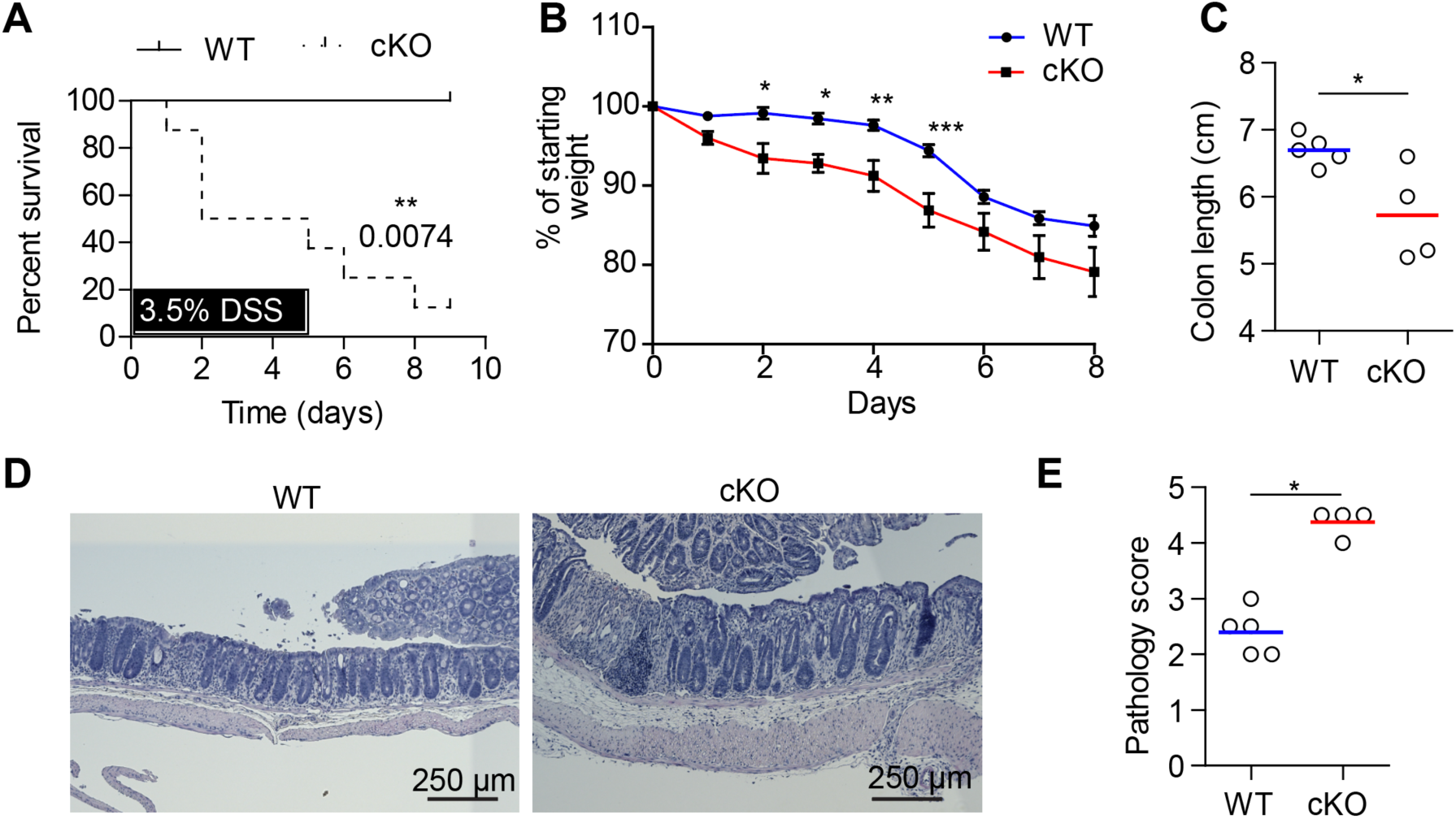
Intestinal-epithelial LSD1 protects mice from chemically induced acute colitis. **A**, Percent survival of male WT and cKO mice after DSS administration (3.5 %) in drinking water for 5 days. Kaplan-Meier survival curves was used and tested with a Gehan-Breslow-Wilcoxon test to identify significant differences between wild-type and cKO after oral DSS administration (P= 0.0074). **B**, Body weight of WT and cKO mice that were administrated DSS in drinking water for 5 days (pooled from 2 experiments). **C**, Colon length of female mice at day 8. **D**, H&E histological analysis of female distal colons of WT and cKO mice at day 8. **E**, Pathology score of DSS-induced colitis in female WT and cKO mice, each dot represents a mouse. Unpaired two-tailed Student’s t test (B, C & E) was performed to observe significant differences. ns = not significant, * P ≤ 0.05 and **, P < 0.01.

### LSD1 is required for defense against *Citrobacter rodentium*

*Citrobacter rodentium*, a gram-negative mucosal bacterium, is used to model the pathogenesis of human enteric pathogens such as Enteropathogenic *Escherichia coli* (Collins et al., 2014). It is currently unknown whether having an epithelial reparative state prior to infection is beneficial for the host. Therefore, we orally infected cKO and WT littermates with *C. rodentium*. Strikingly, we found that cKO mice are highly susceptible to *C. rodentium* with increased mortality and increased weight loss compared to WT littermates (Fig. 3A, 3B). We found a mild increase in the number of *C. rodentium* bacteria in the stool of cKO mice compared to WT mice at day 6 (Fig. 3C). Importantly, at this time point, we already found *C. rodentium* in the liver and spleen of cKO but not in WT mice (Fig. 3D), indicating an early loss of intestinal barrier in these mice. An appropriate immune response is an important aspect in the defense against *C. rodentium*. However, we found near identical responses when assessing cytokine levels of IL-22, IL-17A, and IFN-γ in the colon by qPCR or in re-stimulated mesenteric lymph node cells by ELISA (Fig. 3E, 3F). Thus, although cKO were highly susceptible to infection, their immune response was not atypical.

**Fig. 3.**
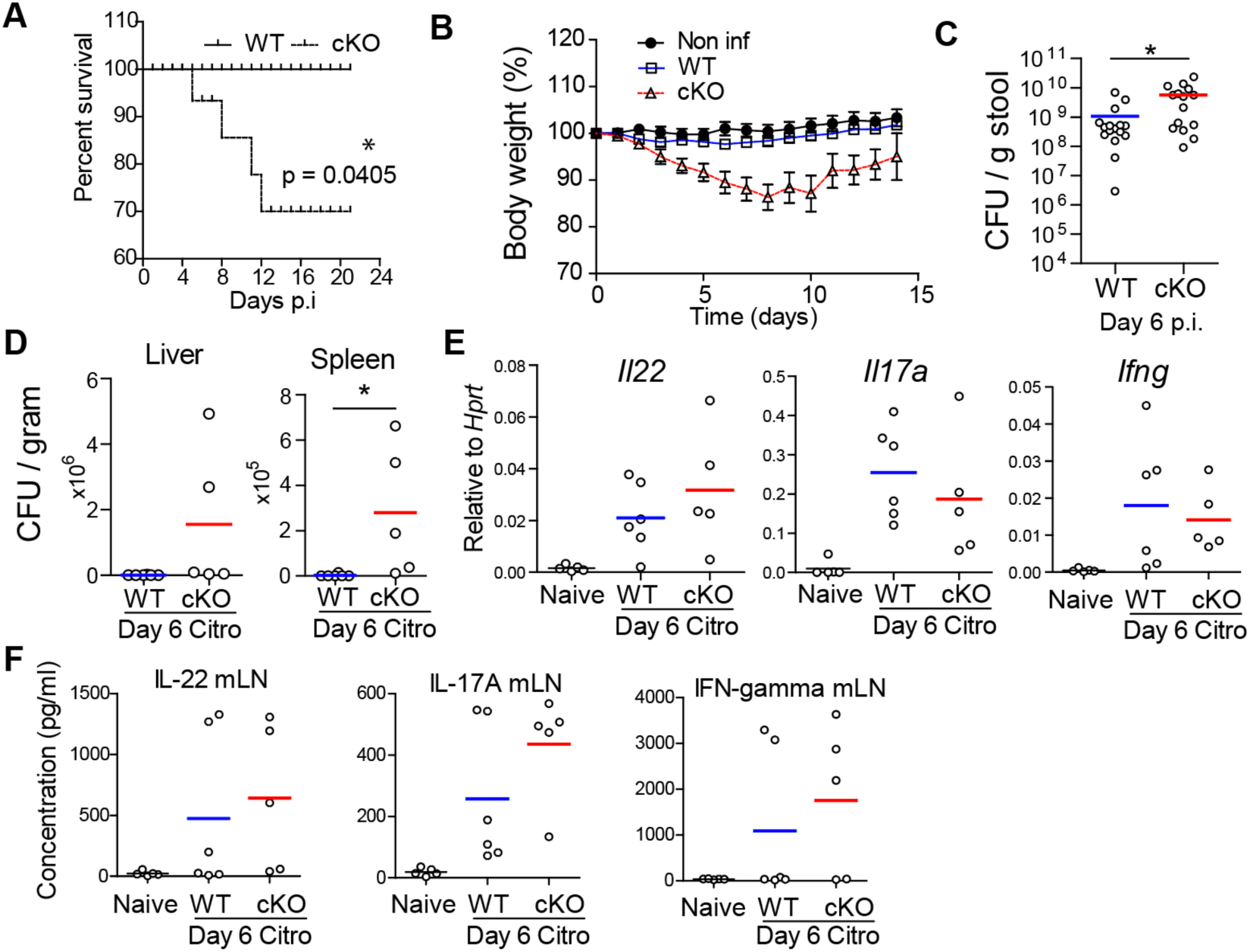
LSD1-deficient epithelium renders mice susceptible to *C. rodentium*. **A**, Survival plot of WT and cKO mice after *C. rodentium* infection. Kaplan-Meier survival curves was used and tested with a Gehan-Breslow-Wilcoxon test to identify significant differences between wild-type and cKO mice (p value= 0.0405). **B**, Body weight loss in Naive, WT and cKO mice during *C. rodentium* infection. n=9 (non inf), n= 16 (WT) and n = 14 (cKO) mice, three pooled experiments. Data point is mean and SD. **C**, CFU (colony forming units) counts from stool using plating on agar. **D**, CFU counts in liver and spleen tissue at day 6 post *C. rodentium* infection. **E**, RT-qPCR for *Il22, Il17a* and *Ifng* was carried out in colon tissue from Naive, WT and cKO mice infected with *C. rodentium* for 6 days. **F**, ELISA for IL-22, IL-17A and IFN-γ in re-stimulated mesenteric lymph node cells after 6 days of *C. rodentium* infection. Each data point represents a mouse, and the mean is depicted. Unpaired two-tailed Student’s t test (C & D) was performed. * P ≤ 0.05. Each dot represents one mouse, and means are depicted.

### LSD1 regulates crypt length and goblet cell responses after *C. rodentium* infection

Next, we examined the epithelial response upon *C. rodentium* infection more closely. We found a slight increase in epithelial-derived antimicrobial *Reg3b* and equal *Reg3g* mRNA levels in the colon comparing cKO with WT mice (Fig. 4A). This indicates that an appropriate immune response and an adequate induction of epithelial-derived antimicrobials in cKO mice is insufficient to prevent morbidity and mortality. It is known that crypt length and induction of goblet cells is essential for resistance against *C. rodentium* and that these are indicators of disease (Bergstrom et al., 2010; Papapietro et al., 2013). Histological studies revealed that there is an increase in colonic crypt hyperplasia in cKO mice compared to WT mice at day 6 post infection (Fig. 4B, 4C), and that these long crypts are characterized by Ki67^+^ cells (Fig. 4D). In addition, by inspection of a dead animal, we found a complete detachment of the epithelium that likely caused the sudden mortality (Fig. S2B). Next, we sought to determine the goblet-cell responses that are induced by *C. rodentium* infection. We found that *C. rodentium* induced goblet-cell effector genes such as *Retnlb* and *Ang4* in WT mice, however, in stark contrast, this induction was completely absent in cKO mice (Fig. 4E), which may contribute to the observed pathology. In summary, although IL-22-induced antimicrobials such as REG3γ are normally regulated, the proliferative and the goblet-cell responses are strongly dependent on epithelial LSD1 expression.

**Fig. 4.**
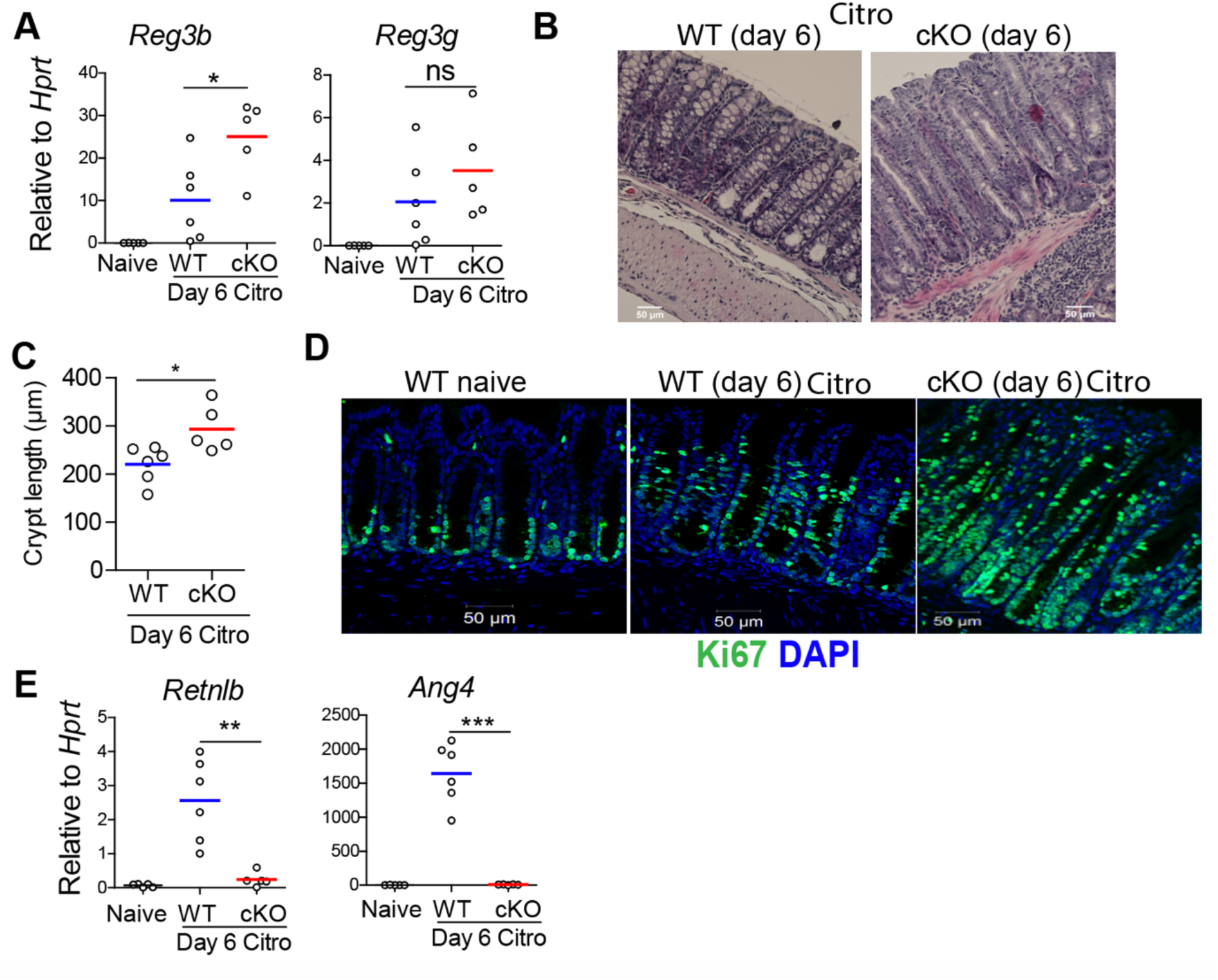
Intestinal-epithelial LSD1 is required for appropriate crypt hyperplasia and goblet cell responses to infection with *C. rodentium*. **A**, RT-qPCR for antimicrobials *Reg3b* and *Reg3g* in colon tissue of Naive (WT), and WT and cKO mice infected with *C. rodentium* for 6 days. **B, C** H&E staining and quantification of crypts length in WT and cKO mice infected with *C. rodentium* for 6 days. **D**, Ki67 staining to measure proliferation in colon tissue of Naive (WT), WT and cKO mice infected with *C. rodentium* for 6 days. **E**, RT-qPCR for goblet-cell specific antimicrobials *Retnlb* and *Ang4* in colon tissue of Naive, WT and cKO mice infected with *C. rodentium* for 6 days. Unpaired two-tailed Student’s t test (A, C & E) was performed to observe significant differences among experimental groups. ns = not significant, * P ≤ 0.05, **, P < 0.01, *** P ≤ 0.001. Each dot represents one mouse and means are depicted.

### LSD1 is required for optimal immunity against the helminth *Trichuris muris*

Resistance to intestinal infection is often an interplay between various cell types in which the epithelium is part of the effector mechanism. *Trichuris muris* is a murine whipworm that is closely related to the human parasite *T. trichiura*. Clearance of *T. muris* is dependent on a type 2 response that leads to increased epithelial turnover, in which the epithelium acts like an escalator to get rid of worms (Cliffe et al., 2005; Oudhoff et al., 2016). In addition, a type 2 response leads to goblet cell hyperplasia, which is also important (Hasnain et al., 2010). To study the role of intestinal-epithelial LSD1 in response to *T. muris*, mice were infected with ∼200 infective embryonated eggs by oral gavage. We observed that parasite elimination was incomplete in cKO mice on day 21 post infection, at which point WT littermates had effectively cleared the infection (Fig. 5A). Goblet cell hyperplasia is a hallmark feature and important for the elimination of worms from the ceacum (Hasnain et al., 2010; Turner et al., 2013). Therefore, we next carried out a histological examination of cKO and WT ceacums at day 21 post *T. muris* infection. Our Periodic acid-Schiff (PAS) staining revealed a near complete absence in goblet cells in the ceacum of *T. muris* infected cKO mice compared to infected WT mice (Fig. 5B, 5C). In support, we also found reduced expression of goblet-cell effector genes *Retnlb* and *Clca1* in infected cKO mice compared to WT mice (Fig. 5D). Elimination of *T. muris* parasites is dependent on expression of type 2 cytokines. Analysis of the hallmark Th2 cytokine revealed that protein expression of IL-13 was almost similar in WT and cKO mice infected with *T. muris* (Fig.5E). However, the expression of the Th1 cytokine IFN-γ was upregulated in restimulated lymph node cells of cKO mice in comparison with WT littermates (Fig. 5E). Therefore, we conclude that upregulated IFN-γ expression in infected cKO mice in combination with a lack of goblet cell responses renders these mice susceptible to *T. muris* infection.

**Fig. 5.**
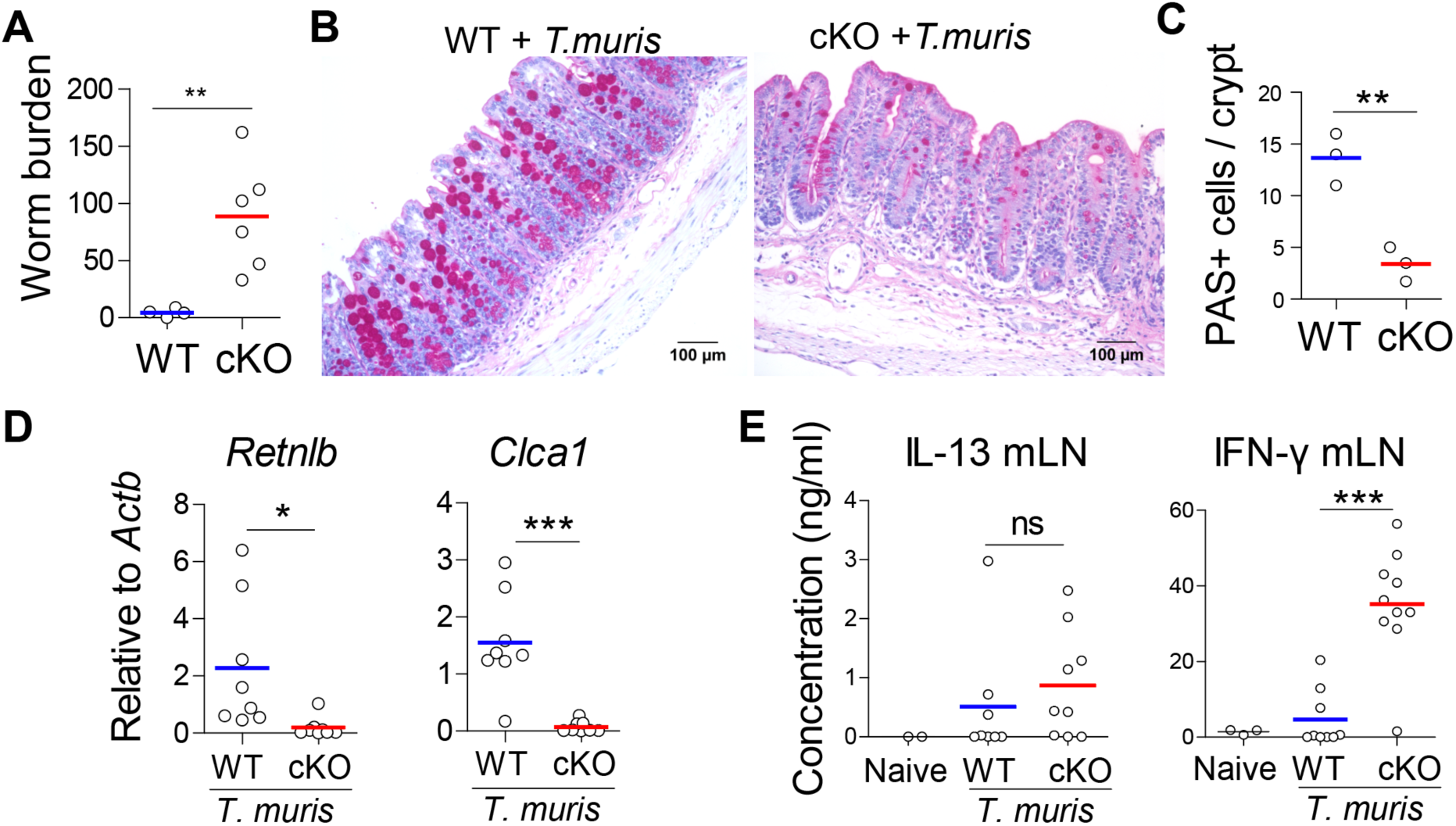
Intestinal-epithelial LSD1 is required for effective immunity against the whipworm *Trichuris muris*. **A**, WT and cKO mice were infected with 200 embryonated eggs of *T. muris*. Worm burden was quantified in caecum after 21 days of infection. Each point is a mouse, pooled from 2 independent experiments. **B**, PAS staining for goblet cells was done in ceacal epithelium of WT and cKO mice after 21 days of *T. muris* infection. **C** Quantification of PAS+ cells in caecal epithelium of WT and cKO mice after 21 days of *T. muris* infection. n=3 from 1 representative experiment. **D**, RT-qPCR for *Retnlb* and *Clca1* in proximal colon of WT and cKO after 21 days of *T. muris* infection. Each data point is 1 mouse. **E**, Sandwich ELISA for quantification of IL-13 and IFN-γ expression in restimulated mesenteric lymph node cells after 21 days of *T. muris* infection. Unpaired two-tailed Student’s t test (A, C D & E) was performed to observe significant differences. ns = not significant, * P ≤ 0.05, **, P < 0.01, *** P ≤ 0.001.

### LSD1 controls homeostatic and cytokine driven goblet cell responses

In both *C. rodentium* and *T. muris* infection models we found a complete inability to induce goblet-cell effector molecules that are normally induced by IL-22 and IL-13 (Fig. 4E, 5D). We have previously found a modest reduction in goblet cell numbers in small intestinal tissues of cKO mice (Zwiggelaar et al., 2020). However, we found that goblet cell differentiation and maturation is more severely impaired in the colons of naïve cKO mice (Fig. 6A-C), and this is similar in GSK-LSD1 treated intestinal organoids (Fig. 6C-E). Indeed, gene set enrichment analysis (GSEA) showed a negative correlation with a goblet cell gene set (Haber et al., 2017) of a previously performed RNA seq experiment of small intestine crypt cells comparing cKO mice with WT littermates (Fig. 6C, sequencing data from (Zwiggelaar et al., 2020)Similarly, the goblet cell gene set was negatively associated with transcriptome analysis of organoids that were treated with GSK-LSD1 for 24 h (Fig. 6C, sequencing data from (Zwiggelaar et al., 2020)). This last part indicates an immediate loss of goblet cell differentiation upon inhibition of LSD1, which is likely independent of H3K4me1/2 enhancer modulation by LSD1 as we would expect this change to be more transient. Instead, it may be due to the disassociation of the LSD1-GFI1 co-repressor complex, which is independent of LSD1 enzyme activity, as was previously suggested and reported (Maiques-Diaz et al., 2018; Zwiggelaar et al., 2020). In support, GFI1 KO mice have a lack of goblet cell differentiation and maturation (Shroyer et al., 2005).

**Fig. 6.**
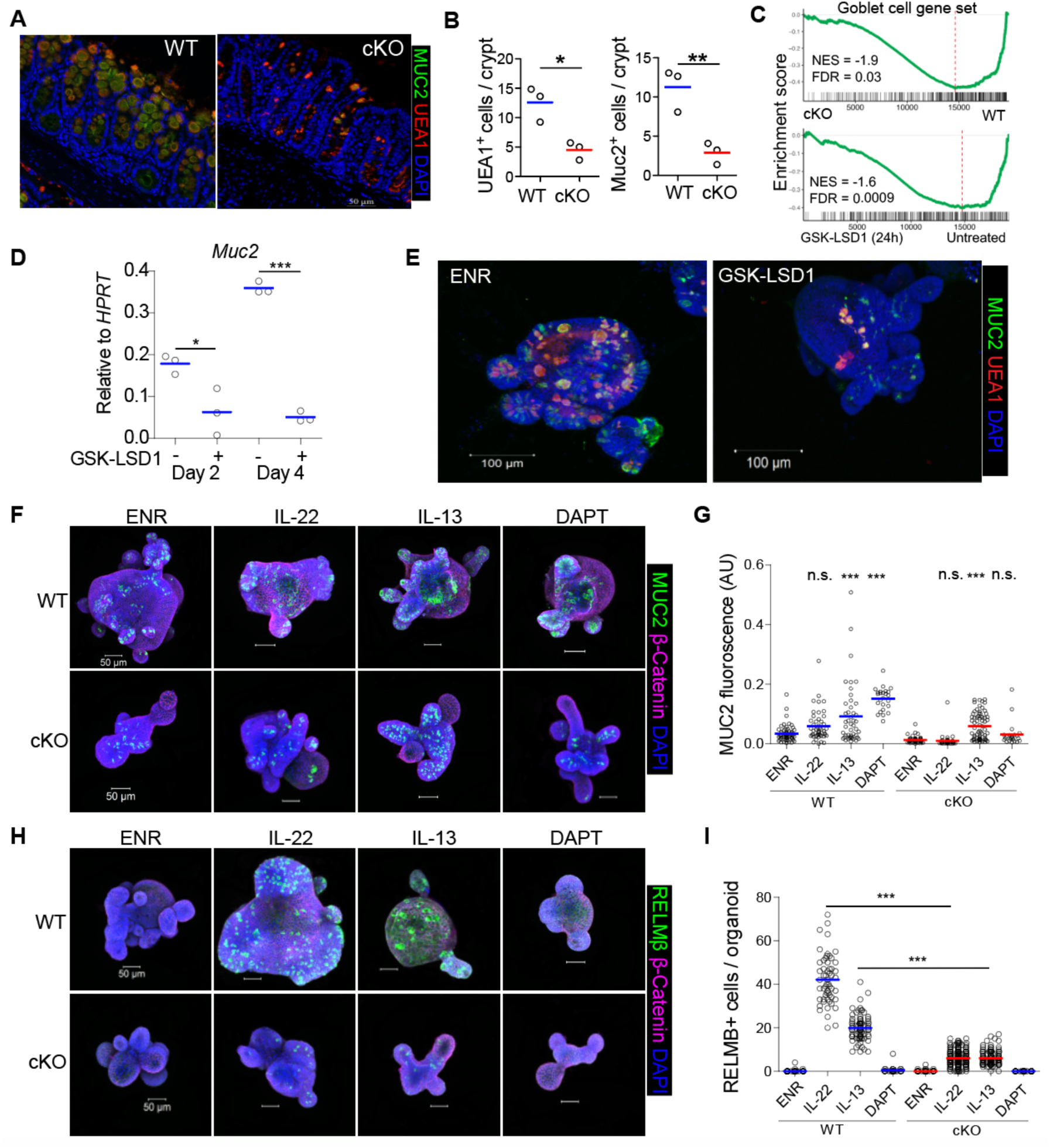
LSD1 controls (immune-driven) goblet cell responses. **A**, Colon of WT and cKO mice were stained for MUC2 (green), Ulex europaeus agglutinin I (UEA1) (red). UEA1 is a lectin which binds to glycoproteins and glycolipids and used as an additional marker for goblet cells. DAPI (blue) used as a counterstaining. **B**, Quantification of MUC2+ cells and UEA1+ cells in the colon of WT and cKO mice. **C**, Application of GSEA of a goblet cell gene set in WT and cKO small intestine RNA seq, and untreated vs GSK-LSD1 (24h) treated organoids RNA-seq data. **D**. RT-qPCR for *Muc2* expression in organoids left untreated or treated with GSK-LSD1 for 2 and 4 days. Expression was normalized to housekeeping gene Hprt. **E**. Immunofluorescent images of MUC2 in WT and GSK-LSD1 treated organoids for 4 days. **F and H**. Immunofluorescence images for MUC2 (F, green) and RELMβ (H, green) in WT and cKO organoids left untreated (ENR) and treated with IL-13 (10 ng/ml), IL-22 (5ng/ml) and DAPT 10 μM for 4 days. DAPI was used to stain nuclei (blue). **G & I**. Quantification of MUC2 fluorescence intensity (G) and number of RELMβ positive cells (I) per organoid. Unpaired two-tailed Student’s t test (B & D) and One-way analysis of variance with Tukey’s Multiple Comparison Test (G & I) was performed to observe significant differences. ns = not significant, * P ≤ 0.05, **, P < 0.01, *** P ≤ 0.001.

To further define goblet cell responses, we treated WT and cKO organoids with IL-13, IL-22, and included DAPT as a positive control. DAPT is a γ-secretase inhibitor that blocks NOTCH signalling to robustly induce goblet cell differentiation. As a readout, we used immunofluorescence staining of MUC2, a canonical goblet cell marker, and RELMβ, a goblet cell effector protein. As we found recently, IL-13 but not IL-22 induces MUC2 in WT organoids (Lindholm et al., 2020), and this response was similar in cKO organoids (Fig. 6F, 6G), suggesting that IL-13 can overrule the lack of goblet cell differentiation caused by LSD1 deficiency. In contrast, DAPT strongly induces MUC2 in WT but not cKO organoids (Fig. 6F, 6G). In addition, DAPT did not induce RELMβ in either WT or cKO cultures (Fig. 6H, 6I), which suggests that the RELMβ effector response does not rely on NOTCH signalling. Nevertheless, both IL-22 and IL-13 caused a dramatic increase in RELMβ+ cells in WT organoids, whereas this was a very subtle response in cKO cultures (Fig. 6H, 6I). To summarize, only IL-13 is able to stimulate MUC2+ goblet cell differentiation in cKO organoids, which occurs independently of NOTCH signalling, and is not sufficient to induce effector responses such as RELMβ accumulation.

### LSD1 controls cell-adhesion and cell-junction gene programs that are important for cell attachment

A lack of goblet cell differentiation and maturation is likely involved in susceptibility to colitis and infection of cKO mice. Similarly, *Muc2* KO mice are susceptible to *C. rodentium* and DSS colitis (Bergstrom et al 2010). However, unlike *Lsd1* cKO mice, *Muc2* KO mice do not mount an increased type 1 response upon *T. muris* infection (Hasnain et al., 2010). In addition, we observed sudden and complete loss of epithelial cells during *C. rodentium* infection and we noticed withering crypts upon DSS colitis (Fig. S2), both of which are not reported in *Muc2* KO mice (Bergstrom et al., 2010). Thus, we hypothesized there may be an additional cause for the pathology we observe in cKO mice upon the different challenges. In our previous work we found that the transcriptomic profile of cKO small intestinal crypt cells is fetal-like (Zwiggelaar et al., 2020). Here, we performed an unbiased analysis of GO terms that are associated with the top 500 upregulated genes in cKO crypt cells compared to WT crypt cells (Fig. 7A). We found several terms associated with the regulation of actin and cytoskeletal processes, which we will discuss below, and we found the enrichment of genes associated with cell-adhesion and cell junction processes (Fig.7A). Tight junctions are crucial in retaining barrier function, and cell adhesion to the basal membrane is important to maintain tissue integrity. Importantly, unlike the goblet cell gene set signature, the genes associated with cell-substrate and cell junction GO terms were not as timely affected upon 24h GSK-LSD1 treatment in organoids (Fig. 7B). Nevertheless, these observations prompted us to check how stable the interaction is between the epithelium and the basal membrane. Therefore, we performed an epithelium cell detachment assay based on EDTA treatment. We observed an increase in detachment of cKO epithelium compared to WT epithelium (Fig. 7C). The ease by which cKO epithelium becomes detached fits well with our observations that large parts of the epithelium were lost during *C. rodentium* infection as well as the withering crypts during DSS colitis. This would also explain the mortality due to sepsis in these colitis models, as well as the induction of a type 1 response upon *T. muris* infection.

**Fig. 7.**
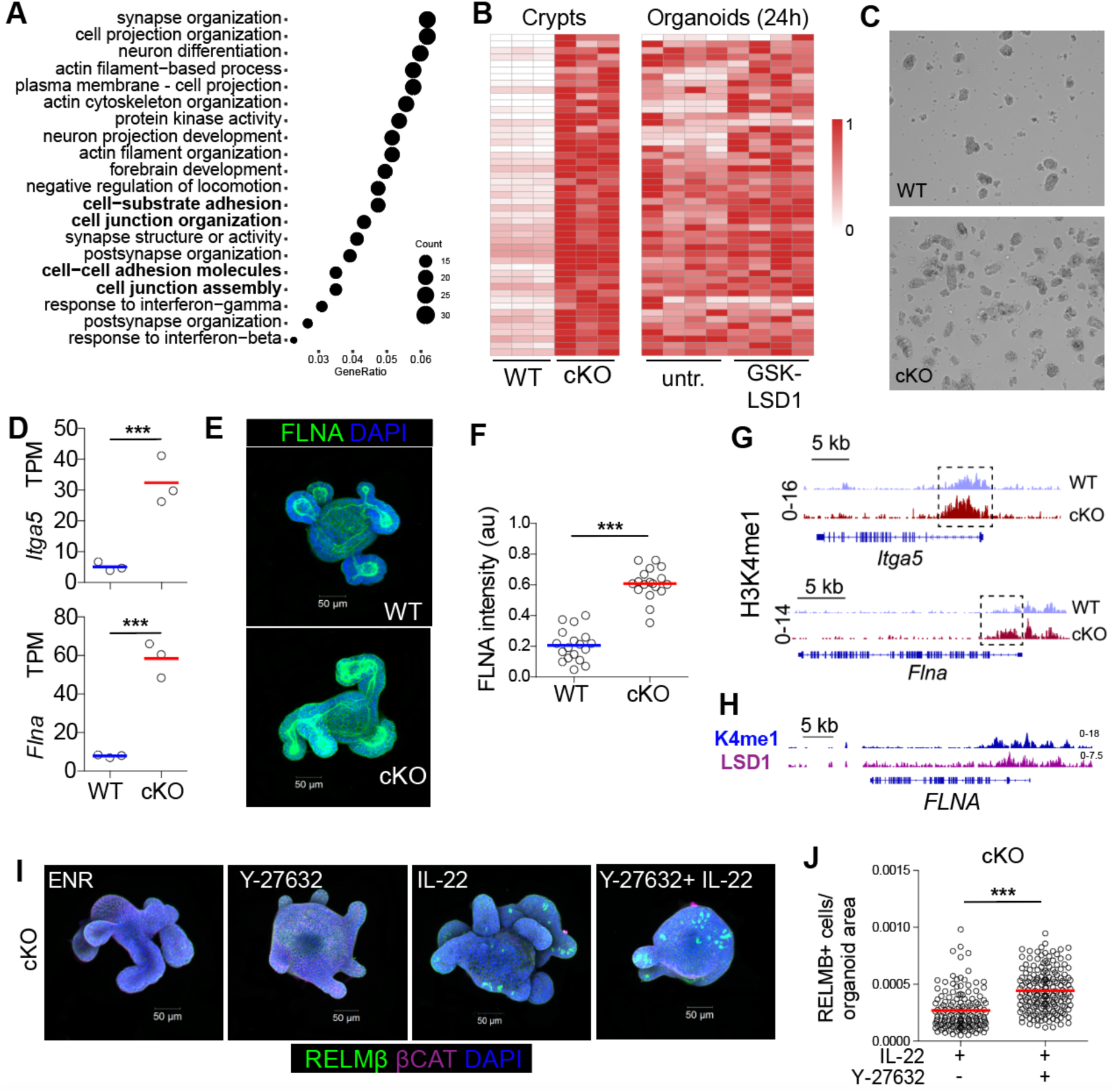
LSD1 controls modulators of the cytoskeleton which is relevant for cell adhesion and goblet cell responses. **A**, Top 500 genes upregulated in cKO crypts were selected and unbiased association with top 20 GO terms is displayed. **B**, Genes taken from in **7A** bold-selected GO terms are displayed in a heatmap using TPM values scaled to the max of the gene comparing WT and cKO crypts (left) and untreated and GSK-LSD1 (24h) treated organoids (right). **C**, Representative images of dissociated epithelial cells from duodenum after EDTA-based cell detachment assay, experiment was performed 3 times with similar results (n=3). **D**, TPM for *Itga5* and *Flna* in WT and cKO small intestine crypts. **E**, Representative confocal images of FILAMIN A (FLNA) stained WT and cKO organoids. **F**, Quantification of FLNA intensity in WT and cKO organoids. **G**, Representative IGV tracks of H3K4me1 levels at *Itga5* and *Flna* genes from WT and cKO crypts. **H**, IGV tracks of H3K4me1 and LSD1 levels at the *FLNA* locus in a human cancer cell line. **I**, cKO organoids were left untreated (ENR) or treated with IL-22 (5ng/ml), Y-27632 (10 μM) and combination of Y-27632 + IL-22 for 4 days and representative images for RELMβ (green) are shown. **J**, Pooled quantification of RELMβ+ cells per organoid area from independent experiments. Unpaired two-tailed Student’s t test (D, F & J) was performed to observe significant differences. ns = not significant, *** P ≤ 0.001.

### LSD1 controls genes associated with cytoskeletal organization to define goblet cell responses

We found that genes associated with cytoskeletal organization were upregulated in cKO crypts (Fig. 7A), and we show *Itga5* and *Flna* as two representative examples (Fig. 7D), We also found increased protein level of FILAMIN A (FLNA) in cKO organoids by confocal microscopy (Fig. 7E, 7F). Importantly, both *Itga5* and *Flna* have increased intragenic H3K4me1 levels in cKO crypts compared to WT crypts (Fig. 7G). In further support, using a published LSD1 ChIP seq data set from a human cancer cell line (Mohammad et al., 2015), we found that LSD1 is enriched at the *FLNA* locus where there is also H3K4me1 (Fig. 7H). This suggests that LSD1, by demethylating H3K4me1 levels at putative enhancer sites, directly controls these important mediators of cell structure and function.

The cytoskeletal network also plays an important role in intestinal epithelial cell differentiation (Petersen et al., 2018). The cytoskeleton can be altered by cytokines, for example, in our recent study we found that IL-13 induces genes associated with the GO term cell shape (Lindholm et al., 2020), and many of these genes were downregulated upon IL-22 treatment (Fig. S3). We hypothesize that IL-13 may rescue MUC2+ goblet cell differentiation in cKO organoids by controlling cell-shape associated genes and thus overrule the restrictions due to LSD1 deficiency. To test if altering the cytoskeleton would also allow for IL-22 mediated goblet cell effector responses, we treated cKO organoids with a combination of IL-22 and Y-27632, an inhibitor of ROCK signalling that is commonly used to manipulate the cytoskeletal network. Indeed, Y-27632 enhanced RELMβ expression in IL-22 treated cKO organoids compared to IL-22 treatment alone (Fig. 7I and J). In summary, we provide a framework in which LSD1 putatively controls enhancers of genes that shape the cytoskeletal organization of intestinal epithelial cells, which is needed for both cell attachment and cytokine-driven goblet cell effector responses after infection.

## DISCUSSION

The gastrointestinal system is continuously exposed to various pathogens, chemicals, and dietary components that can result in inflammation and tissue damage. It was previously found that the epithelium, after damage or infection, temporarily transitions into a reparative fetal-like state that is necessary to regain homeostasis (Nusse et al., 2018; Yui et al., 2018). Thus, drugs that would induce this transition of the epithelium could be attractive therapeutics used in patients with infection or inflammatory bowel disease to aid in the tissue repair, or even prevent disease. Indeed, in a previous report, we showed that reparative epithelium is beneficial upon irradiation injury and that LSD1 is a druggable target that could facilitate the reprogramming (Zwiggelaar et al., 2020). In addition, treating mice with GSK-LSD1 upon damage alleviates colitis symptoms in DSS-colitis (Oh et al., 2020). In this study, we tested whether a pre-existing epithelial reparative state, such as in LSD1 cKO mice, would be beneficial in colitis or infectious disease models. In contrast to previous work, we found that LSD1 cKO mice are susceptible to both inflammatory and infectious disease models. One of the differences is that in our current work the disease models are likely all affected by the pre-existing cytoskeletal features affecting the intestinal epithelial cells. Indeed, we find clear evidence that goblet cell effector genes rely on LSD1, both under homeostasis and upon induction in the diverse disease models. In addition, using an unbiased analysis of previous RNA-seq data, we found that genes associated with modulation of the cytoskeleton including those involved in cell adhesion and tight junctions are regulated by LSD1 (Fig. 7). As an example, we find that the gene encoding for FILAMIN A, a key protein in mechanosensing and mechanotransduction (Zhou et al., 2010), is a direct target of LSD1 demethylase activity (Fig. 7D-H). This could also be a connection of how LSD1 mediates genes that are controlled by the mechanosensing regulatory proteins YAP/TAZ (Zwiggelaar et al., 2020).

LSD1 was the first histone demethylase discovered and has been primarily studied in developmental biology and cancer (Cao et al., 2017; Chen et al., 2017; Wasson et al., 2016). Of note, it is currently a target in several clinical trials for the treatment of cancers. Our data suggests that these patients may experience higher susceptibility to gastrointestinal insults. Nevertheless, the mode of action of LSD1 in the various systems is not always clear. For example, it was found that inhibition of LSD1 leads to differentiation of myeloid leukemia cells, but that this was due to the scaffolding function of LSD1, rather than its histone demethylase activity (Maiques-Diaz et al., 2018). We propose a similar mechanism to be responsible for the immediate inhibition of goblet cell differentiation, in which the GFI1-LSD1 co-repressor complex may be rapidly disrupted (Fig. 6). However, the classical mode of action by LSD1 is demethylation of enhancer regions (Mendenhall et al., 2013). The kinetics that would affect these genes may be a bit slower, and this is indeed what we observe in the regulation of cytoskeletal-modulation associated genes such as *Itga5* and *Flna* (Fig. 7). We propose that this latter mechanism is important in the control of cell attachment as well as goblet cell effector responses such as those induced by IL-13 and IL-22. Together, we propose that the control by LSD1 of genes involved in the cytoskeletal organization is critical for intestinal epithelial responses to inflammation and infection.

## ACKNOWLEDGEMENTS

We thank Drs. Stuart Orkin and Sylvie Robine for kindly sharing mouse strains. We thank the imaging (CMIC) and animal care (CoMed) core facilities that assisted in this work (NTNU). Funding of this work was provided by the Norwegian Research Council (Centre of Excellence grant 223255/F50, and ‘Young Research Talent’ 274760 to MJO) and the Norwegian Cancer Society (182767 to MJO). MMA is the recipient of a Marie Skłodowska-Curie IF (DLV-794391). This work was also supported by the South-Eastern Norway Regional Health Authority, Early Career Grant 2016058, and the Research Council of Norway “Young Research Talent” grant to JAD. MF is supported by the Norwegian Research Council (grant no. 275286). BAV is the CH.I.L.D. Foundation Chair in Pediatric Gastroenterology.

## EXPERIMENTAL PROCEDURES

### Mouse strains

*Villin-Cre* were a kind gift from Sylvie Robine (Marjou et al., 2004) and *Lsd1*^f/f^ mice were a kind gift from Dr. Stuart Orkin (Kerenyi et al., 2013). Mice breeding was carried out and were housed at the Comparative Medicine Core Facility (CoMed).

### *Citrobacter rodentium* infection and CFU counts

*Citrobacter rodentium* infection studies were performed following the guidelines and recommendations for the care and use of animals in research and were approved by Norwegian Food Safety Authority (FOTS ID: 11842). Briefly, GFP-expressing *C. rodentium* (Bergstrom et al., 2010) was grown at 37°C in Luria-Bertani (LB) medium supplemented with chloramphenicol (30 μg/ml). 6-10 weeks old mice were infected by oral gavage with overnight culture of 10^8^ – 10^9^ CFU per mouse delivered in a volume of 0.1ml sterile PBS. Mice were daily monitored after the oral administration with *C. rodentium* and assessed for weight loss and other pain associated behaviors like hunchbacked posture, rectal prolapse and piloerection. Fecal samples were collected, weighed and homogenized in a sterile PBS using FastPrep homogenizer from MP biomedicals. Serially diluted homogenates were plated on chloramphenicol resistant agar plates and number of colonies were counted after incubation of plates at 37°C for 18-24 h. At the end of every experiment, mice were humanely euthanized by inhalation of CO2.

### *Trichuris muris* infection in mice

All animal procedures were performed at the University of British Columbia following institutional guidelines. Briefly, mice were infected by oral gavage with high dose of approximated 200 embryonated *<u>T. muris</u>* eggs. After 21 days of *T. muris* infections, mice were sacrificed for the analysis of worm burden in the ceacum.

### DSS (Dextran sulfate sodium) colitis model in mice

DSS induced colitis was performed following the guidelines and recommendations for the care and use of animals in research and were approved by Norwegian Food Safety Authority (FOTS ID: 11842). Colitis was induced in WT and cKO mice by the administration of Dextran Sulfate Sodium Salt (36,000-50,000.) colitis Grade at a 3.5 %, followed by 3 days of rest. All the mice were daily monitored for weight loss and other signs of illness such as rectal prolapse or blood in the stool. After the completion of the experiment (day 8), mice were euthanized by humane procedures by using CO2 inhalation. Injury score was evaluated by two different researchers in a genotype-blinded way. Features of damage in colon were evaluated in accordance to the level of tissue damage, immune infiltrate presence and loss of cell types/crypts as previously described (Erben, 2014).

### Immunofluorescence and immunohistochemistry staining

After euthanization of mice, segments of intestinal mouse were dissected and washed with sterile PBS and then rolled as swiss roll. Mice intestinal tissue were fixed with 4 % formaldehyde for 48 h at room temperature. Tissue was subsequently dehydrated through a series of ethanol graded and then embedded in the paraffin wax blocks. 5 μm thick sections were prepared using microtome and tissue was placed on glass slides. The tissue slides were then deparaffinized and rehydrated, followed by antigen retrieval. Sections were blocked in blocking buffer (1 % BSA and 2% normal goat serum) for 1 h at room temperature in a humidified chamber. Tissue were incubated with diluted primary antibody (LSD1, 1: 200 Cell Signaling Technology 2184), MUC2 1: 200 Santa Cruz Biotechnology sc-15334), anti Ki67, 1:200 Invitrogen MA5-14520) for overnight at 4°C. Next day, slides were washed with 0.2% PBST for 10mins each wash. Tissue was incubated with secondary antibody coupled to fluorochromes, UEA-1 (1:500 Vector Laboratories RL-1062) and counterstained with DAPI for 1 h at room temperature in the dark. After incubation, slides were washed 3 times with 0.2% PBST for 10 mins each and then mounted in Fluoromount G. Hematoxylin and Eosin (H&E) staining and Periodic Acid Schiff (PAS) staining in paraffin sections was performed according to the standard histological procedures.

### RNA isolation and qPCR

Total RNA was isolated from organoids and colon tissue using the Quick-RNA™ MicroPrep Kit from Zymo Research with slight modifications. Briefly, organoids were first lysed in RNA-Solv Reagent provided by Omega Biotek. For RNA isolation from colon tissue, small piece of colon tissue was first homogenized using FastPrep homogenizer and was resuspended in RNA-Solv Reagent. After lysis, cell lysates were centrifuged at ≥12,000 x g for 1 minute for the removal of particulate and supernatant was transferred to a fresh tube. Equal volume of ethanol (95-100%) was added to the sample and mixture was centrifuged in Zymo-Spin™ IC Column. DNase treatment was performed by adding 80μl of DNase mix directly on to the column and incubated for 15 minutes. To wash the column, 400 µl RNA Prep Buffer to the column and centrifuge for 30 seconds and followed by 700 µl RNA Wash Buffer to the column and centrifuge for 30 seconds. To ensure complete removal of wash buffer, 400 µl RNA of Wash Buffer was added and column was centrifuged for 2 minutes at 12,000 x g. RNA was eluted into the fresh tube in a volume of 20 μL for organoids and 40 μL for colon tissue by addition of DNase/RNase-Free water onto the column. The eluted RNA was used immediately or stored at −80°C for downstream use. Equal amount of purified RNA was reversed transcribed to cDNA using the Applied Biosystem High-Capacity RNA-to-cDNA Kit. Quantitative real-time PCR was performed using the StepOnePlus Real-Time PCR System by using both TaqMan Universal PCR Master Mix and SYBR Green PCR Master Mix by Applied Biosystem.

### RNA Seq analysis

RNA sequencing data from intestinal epithelium with conditional LSD1 KO (acession E-MTAB-7862) and small intestinal organoids treated with GSK-LSD1 (accession E-MTAB-9077) were analyzed as described previously(Zwiggelaar et al., 2020). GSEA analysis was run with log2fold change as weights, 10000 permutations and otherwise default settings using the R-package clusterProfiler(Yu et al., 2012). The clusterProfiler package was also used for GO-term analysis for terms in biological process. Heatmaps were created using the R-package pheatmap.

### ChIP Seq analysis

H3K4me1 ChIP sequencing data from intestinal epithelium with conditional LSD1 KO was analyzed as described previously (Zwiggelaar et al., 2020)(accession E-MTAB-7871) and processed bigwig files from LSD1 ChIP sequencing where downloaded from accession GSE66298 (Mohammad et al., 2015). Plots of ChIP seq signal where made using integrative genomics viewer from bigwig files (James T Robinson et al., 2011).

### Organoids IF staining

For immunofluorescence assays, organoids were seeded in pre-warmed 8 well chamber IBIDI slides. After incubation, organoids were fixed in 4 % paraformaldehyde with 2% sucrose for 30 minutes. Wells with fixed organoids were washed twice with sterile PBS for 5 mins each. Next, organoids were permeabilized with 0.2% Triton X 100 prepared in PBS and incubated for 30 min at room temperature. To block free aldehydes groups and prevent high background signal, organoids were incubated with 100mM glycine for 1 h at room temperature. The organoids were then incubated in the blocking buffer (1% BSA, 2 % NGS diluted in 0.2% Triton X 100 in PBS) in a humidified chamber for 1 h at room temperature. Organoids were incubated in the diluted primary antibody in the same blocking buffer for overnight at 4°C. Next day, the primary antibody anti ki67(1:200 abcam ab15580) Phospho-Stat3 (1:100, Cell Signaling Technology, 9145), STAT3 (1:1600, Cell Signaling Technology 9139) MUC2 (1:200, Santa Cruz Biotechnology sc-15334), β-catenin 1:200, Santa Cruz Biotechnology sc-7963), Filamin A (1:200, Invitrogen PA5-82043), RELMβ (1:200, Peprotech 500-p215) solution was decanted, and wells were washed three times for 10 minutes each with PBS with slight agitation. Organoids were incubated with secondary antibody(1:500, Goat anti-Rabbit IgG Alexa Fluor 488 Invitrogen A-11034 and 1:500, Goat anti-Mouse IgG Alexa Fluor 647 Invitrogen A-21236), Ulex europaeus agglutinin-1 (UEA1, 1:500 Vector Laboratories RL-1062 and DAPI (1:1000) as counterstained for overnight at 4°C in the dark. Next day, secondary antibody solution was decanted, and wells were washed three times with PBS for 10 minutes in agitation. After staining’s, wells were mounted in 250 μL of Fluoromount G and were visualized with a confocal microscope.

### Western blot analysis

Total protein was extracted from the organoids. Briefly, organoids were harvested and resuspended in RIPA Lysis and Extraction Buffer supplemented with PhosSTOP Phosphatase Inhibitor Cocktail for 10mins in ice. Proteins (50μg) were separated by NuPAGE Bis-Tris protein precast polyacrylamide gels and transferred to nitrocellulose membranes using iBlot 2 Gel Transfer device. The membranes were incubated with 5 % blocking buffer (5 % BSA in TBST) for 1h at room temperature. The membranes were further incubated with the primary antibodies against p-STAT3(1:2000, Cell Signaling Technology, 9145), STAT3(1:1000, Cell Signaling Technology 9139) and GAPDH (1:5000, abcam ab125247) for overnight at 4°C. Next day, after three washes with TBST for 5 mins, membranes were incubated with secondary antibody conjugated with horseradish peroxidase for 1 h at room temperature. Again, membranes were washed three times with TBST for 5 minutes and were visualized with Super Signal West Femto Maximum Sensitivity Substrate and images were acquired using Licor Odyssey detection system.

### Mesenteric lymph nodes isolation and CD3/CD28 restimulation

The mesenteric lymph nodes were isolated from naïve or infected mice and kept in sterile PBS. Harvested lymph nodes from individual mouse were passed through 70 μm cell strainer for single cell suspension. The cell viability was assessed using Trypan blue and counted using Countess™ automated cell counter (Invitrogen). The cells were resuspended in DMEM and adjusted to 2×10^5^ cells per well. A solution of anti-mouse CD3e (100314, Biolegend) was prepared at 5 µg/mL in sterile PBS and 50 μl of antibody solution was added into 96 well plate followed by incubation at 37°C for 2 hours. The antibody solution was removed with a multichannel pipettor and single cell suspension of lymph node cells (2×10^5^ cells per well) was added to respective wells in coated 96 well plates. Next, soluble anti-mouse CD28 (102112, Biolegend) was added to cells at 2 µg/mL. Cells were incubated for 4 days to induce proliferation of cells. After incubation, cells were centrifuged and supernatant was collected for cytokine analysis by ELISA.

### Flow cytometry

Colon from WT and cKO mice were dissected and flushed by PBS. Colon was open longitudinally and cut into small piece of fragments. These fragments were further washed with cold PBS until the supernatant was clear. Next, the colon fragments were incubated with crypts isolation buffer (20mM EDTA in PBS) for 45 minutes at cold room with slight agitation. After incubation, EDTA buffer was removed and fragments were vigorously resuspended in 20ml of ice chilled PBS by shaking and 10mL pipette. The supernatants with crypts was collected in 50ml falcon tube coated with BSA. Next, supernatants with crypts was centrifuged at 300 g for 5 minutes at 4°C to pellet down the crypts. Crypts were further resuspended in TryplE express for 30 minutes at 37°C and were dissociated with pipette every 5 minutes. Digested crypts cells were incubated with anti-mouse TruStain Fc (1:400, BioLegend 101320) for 5 minutes in ice. Cells were further stained with Alexa Fluor 488 anti-mouse SCA1 antibody (1:200, BioLegend 108116) and other epithelial cells markers (anti-mouse CD326 Brilliant Violet 605, 1:200 Biolegend 118227 and PerCP/Cy5.5 anti-mouse CD24, 1:200 Biolegend 101824) for 20 minutes at 4°C, twice washed with PBS and incubated with DAPI for 3 minutes in the last step. Before acquisition, cells suspension was passed through a 70 µm filter mesh. All the samples were acquired using a BD LSR II flow cytometer (BD Biosciences) and analyzed using a FlowJo software.

### Epithelial cell detachment assay

The epithelial cell detachment assay was performed as published with minor modifications (De Arcangelis et al., 2017). The duodenum from WT and cKO mice was dissected and flushed by using PBS. The duodenal tissue was weighed and then opened longitudinally and cut into very small pieces. Importantly, the duodenum was not scraped during this process. Next, the fragments were washed with PBS for equal number of times until the supernatant becomes fully clear. After washing, these small fragments were incubated with different EDTA concentration sequentially (0.5 mM, 2mM and 10 mM) in PBS for 15 mins each at 4°C with slight rotation. The dissociated cells during the rotation were collected by centrifugation at 300 g for 6 mins. These dissociated cells were imaged using EVOS 2 after EDTA incubation.

### Statistical analysis

The statistical analysis was performed using the Graph Pad Prism. Unpaired two-tailed Student’s t test and One-way analysis of variance with Tukey’s Multiple Comparison Test were used to determine the Statistically significant differences between the different experimental groups and P value of <0.05 was considered significant. All the Statistical tests used are described in the figure legends.

## REFERENCES

Allaire, J.M., Crowley, S.M., Law, H.T., Chang, S.Y., Ko, H.J., and Vallance, B.A. (2018). The Intestinal Epithelium: Central Coordinator of Mucosal Immunity. Trends Immunol. 39, 677–696.

De Arcangelis, A., Hamade, H., Alpy, F., Normand, S., Bruyère, E., Lefebvre, O., Méchine-Neuville, A., Siebert, S., Pfister, V., Lepage, P., et al. (2017). Hemidesmosome integrity protects the colon against colitis and colorectal cancer. Gut 66, 1748–1760.

Benz, H.L., Ehman, E.C., Johnson, G.B., Villanueva-meyer, J.E., Cha, S., Leynes, A.P., Eric, P., Larson, Z., and Hope, T.A. (2017). Wound repair: role of immune–epithelial interactions. 46, 1247–1262.

Bergstrom, K.S.B., Kissoon-Singh, V., Gibson, D.L., Ma, C., Montero, M., Sham, H.P., Ryz, N., Huang, T., Velcich, A., Finlay, B.B., et al. (2010). Muc2 protects against lethal infectious colitis by disassociating pathogenic and commensal bacteria from the colonic mucosa. PLoS Pathog. 6.

Bevins, C.L., and Salzman, N.H. (2011). Paneth cells, antimicrobial peptides and maintenance of intestinal homeostasis. Nat. Rev. Microbiol. 9, 356–368.

Biton, M., Haber, A.L., Rogel, N., Burgin, G., Beyaz, S., Schnell, A., Ashenberg, O., Su, C.-W., Smillie, C., Shekhar, K., et al. (2018). T Helper Cell Cytokines Modulate Intestinal Stem Cell Renewal and Differentiation. Cell 175, 1307-1320.e22.

Cai, J., Zhang, N., Zheng, Y., De Wilde, R.F., Maitra, A., and Pan, D. (2010). The Hippo signaling pathway restricts the oncogenic potential of an intestinal regeneration program. Genes Dev. 24, 2383–2388.

Cao, C., Vasilatos, S.N., Bhargava, R., Fine, J.L., Oesterreich, S., Davidson, N.E., and Huang, Y. (2017). Functional interaction of histone deacetylase 5 (HDAC5) and lysine-specific demethylase 1 (LSD1) promotes breast cancer progression. Oncogene 36, 133–145.

Carlet, J. (2012). The gut is the epicentre of antibiotic resistance. Antimicrob. Resist. Infect. Control 1, 1.

Chen, J., Ding, J., Wang, Z., Zhu, J., Wang, X., and Du, J. (2017). Identification of downstream metastasis-associated target genes regulated by LSD1 in colon cancer cells. Oncotarget 8, 19609–19630.

Chiacchiera, F., Rossi, A., Jammula, S., Zanotti, M., and Pasini, D. (2016). PRC 2 preserves intestinal progenitors and restricts secretory lineage commitment. EMBO J. 35, 2301–2314.

Cliffe, L.J., Humphreys, N.E., Lane, T.E., Potten, C.S., Booth, C., and Grencis, R.K. (2005). Immunology - Accelerated intestinal epithelial cell turnover: A new mechanism of parasite expulsion. Science (80-.). 308, 1463–1465.

Collins, J.W., Keeney, K.M., Crepin, V.F., Rathinam, V.A.K., Fitzgerald, K.A., Finlay, B.B., and Frankel, G. (2014). Citrobacter rodentium: infection, inflammation and the microbiota. Nat. Publ. Gr. 12, 612–623.

Erben, et al (2014). A guide to histomorphological evaluation of intestinal inflammation in mouse models. Int. J. Clin. Exp. Pathol. 7, 4557–4576.

Haber, A.L., Biton, M., Rogel, N., Herbst, R.H., Shekhar, K., Smillie, C., Burgin, G., Delorey, T.M., Howitt, M.R., Katz, Y., et al. (2017). A single-cell survey of the small intestinal epithelium. Nature 551, 333–339.

Hasnain, S.Z., Wang, H., Ghia, J.E., Haq, N., Deng, Y., Velcich, A., Grencis, R.K., Thornton, D.J., and Khan, W.I. (2010). Mucin Gene Deficiency in Mice Impairs Host Resistance to an Enteric Parasitic Infection. Gastroenterology 138, 1763-1771.e5.

Jadhav, U. (2017). Acquired tissue-specific promoter bivalency is a basis for PRC2 necessity in adult cells. Physiol. Behav. 176, 139–148.

James T Robinson, Thorvaldsdóttir, H., Winckler, W., Guttman, M., Lander, E.S., Getz, G., and Mesirov, J.P. (2011). Integrative genomics viewer. Nat. Biotechnol. 29, 24–26.

Kerenyi, M.A., Shao, Z., Hsu, Y.J., Guo, G., Luc, S., O’Brien, K., Fujiwara, Y., Peng, C., Nguyen, M., and Orkin, S.H. (2013). Histone demethylase Lsd1 represses hematopoietic stem and progenitor cell signatures during blood cell maturation. Elife 2013, 1–23.

Kim, Y.S., and Ho, S.B. (2010). Intestinal goblet cells and mucins in health and disease: Recent insights and progress. Curr. Gastroenterol. Rep. 12, 319–330.

Kirk, M.D., Pires, S.M., Black, R.E., Caipo, M., Crump, J.A., Devleesschauwer, B., Döpfer, D., Fazil, A., Fischer-Walker, C.L., Hald, T., et al. (2015). World Health Organization Estimates of the Global and Regional Disease Burden of 22 Foodborne Bacterial, Protozoal, and Viral Diseases, 2010: A Data Synthesis. PLoS Med. 12, e1001921.

Koppens, M.A.J., Bounova, G., Gargiulo, G., Tanger, E., Janssen, H., Cornelissen-Steijger, P., Blom, M., Song, J.Y., Wessels, L.F.A., and van Lohuizen, M. (2016). Deletion of Polycomb Repressive Complex 2 From Mouse Intestine Causes Loss of Stem Cells. Gastroenterology 151, 684-697.e12.

Kouzarides, T. (2007). Chromatin modifications and their function. Cell 128, 693–705.

Lindemans, C.A., Calafiore, M., Mertelsmann, A.M., Margaret, H., Connor, O., Dudakov, J.A., Jenq, R.R., Velardi, E., Young, L.F., Smith, O.M., et al. (2010). Interleukin-22 Promotes Intestinal Stem Cell-Mediated Epithelial Regeneration. Nature 528, 169.

Lindholm, H.T., Parmar, N., Drurey, C., Ostrop, J., Díez-Sanchez, A., Maizels, R.M., and Oudhoff, M.J. (2020). Developmental pathways regulate cytokine-driven effector and feedback responses in the intestinal epithelium. BioRxiv 2020.06.19.160747.

Maiques-Diaz, A., Spencer, G.J., Lynch, J.T., Ciceri, F., Williams, E.L., Amaral, F.M.R., Wiseman, D.H., Harris, W.J., Li, Y., Sahoo, S., et al. (2018). Enhancer Activation by Pharmacologic Displacement of LSD1 from GFI1 Induces Differentiation in Acute Myeloid Leukemia. Cell Rep. 22, 3641–3659.

Marjou, F., Janssen, K.P., Chang, B.H.J., Li, M., Hindie, V., Chan, L., Louvard, D., Chambon, P., Metzger, D., and Robine, S. (2004). Tissue-specific and inducible Cre-mediated recombination in the gut epithelium. Genesis 39, 186–193.

Mendenhall, E.M., Williamson, K.E., Reyon, D., Zou, J.Y., Ram, O., Joung, J.K., and Bernstein, B.E. (2013). Locus-specific editing of histone modifications at endogenous enhancers. Nat. Biotechnol. 31, 1133–1136.

Mohammad, H.P., Smitheman, K.N., Kamat, C.D., Soong, D., Federowicz, K.E., Van Aller, G.S., Schneck, J.L., Carson, J.D., Liu, Y., Butticello, M., et al. (2015). A DNA Hypomethylation Signature Predicts Antitumor Activity of LSD1 Inhibitors in SCLC. Cancer Cell 28, 57–69.

Nusse, Y.M., Savage, A.K., Marangoni, P., Rosendahl-Huber, A.K.M., Landman, T.A., De Sauvage, F.J., Locksley, R.M., and Klein, O.D. (2018). Parasitic helminths induce fetal-like reversion in the intestinal stem cell niche. Nature 559, 109–113.

Oh, C., Jeong, J., Oh, S.K., Baek, S.H., and Kim, K. Il (2020). Inhibition of LSD1 phosphorylation alleviates colitis symptoms induced by dextran sulfate sodium. BMB Rep.

Oudhoff, M.J., Antignano, F., Chenery, A.L., Burrows, K., Redpath, S.A., Braam, M.J., Perona-Wright, G., and Zaph, C. (2016). Intestinal Epithelial Cell-Intrinsic Deletion of Setd7 Identifies Role for Developmental Pathways in Immunity to Helminth Infection. PLoS Pathog. 12, 1–16.

Papapietro, O., Teatero, S., Thanabalasuriar, A., Yuki, K.E., Diez, E., Zhu, L., Kang, E., Dhillon, S., Muise, A.M., Durocher, Y., et al. (2013). R-Spondin 2 signalling mediates susceptibility to fatal infectious diarrhoea. Nat. Commun. 4.

Petersen, N., Frimurer, T.M., Terndrup Pedersen, M., Egerod, K.L., Wewer Albrechtsen, N.J., Holst, J.J., Grapin-Botton, A., Jensen, K.B., and Schwartz, T.W. (2018). Inhibiting RHOA Signaling in Mice Increases Glucose Tolerance and Numbers of Enteroendocrine and Other Secretory Cells in the Intestine. Gastroenterology 155, 1164-1176.e2.

Rubin, D.C., Shaker, A., and Levin, M.S. (2012). Chronic intestinal inflammation: Inflammatory bowel disease and colitis-associated colon cancer. Front. Immunol. 3, 1–10.

Shi, Y., Lan, F., Matson, C., Mulligan, P., Whetstine, J.R., Cole, P.A., Casero, R.A., and Shi, Y. (2004). Histone demethylation mediated by the nuclear amine oxidase homolog LSD1. Cell 119, 941–953.

Shroyer, N.F., Wallis, D., Venken, K.J.T., Bellen, H.J., and Zoghbi, H.Y. (2005). Gfi1 functions downstream of Math1 to control intestinal secretory cell subtype allocation and differentiation. Genes Dev. 19, 2412–2417.

Ting, H.-A., and von Moltke, J. (2019). The Immune Function of Tuft Cells at Gut Mucosal Surfaces and Beyond. J. Immunol. 202, 1321–1329.

Turner, J.E., Stockinger, B., and Helmby, H. (2013). IL-22 Mediates Goblet Cell Hyperplasia and Worm Expulsion in Intestinal Helminth Infection. PLoS Pathog. 9, 1–7.

Wang, Y., Chiang, I.-L., Ohara, T.E., Fujii, S., Cheng, J., Muegge, B.D., Ver Heul, A., Han, N.D., Lu, Q., Xiong, S., et al. (2019). Long-Term Culture Captures Injury-Repair Cycles of Colonic Stem Cells. Cell 179, 1144-1159.e15.

Wasson, J.A., Simon, A.K., Myrick, D.A., Wolf, G., Driscoll, S., Pfaff, S.L., Macfarlan, T.S., and Katz, D.J. (2016). Maternally provided LSD1/KDM1A enables the maternal-to-zygotic transition and prevents defects that manifest postnatally. Elife 5, 1–25.

Yu, G., Wang, L.G., Han, Y., and He, Q.Y. (2012). ClusterProfiler: An R package for comparing biological themes among gene clusters. Omi. A J. Integr. Biol. 16, 284–287.

Yui, S., Azzolin, L., Maimets, M., Pedersen, M.T., Fordham, R.P., Hansen, S.L., Larsen, H.L., Guiu, J., Alves, M.R.P., Rundsten, C.F., et al. (2018). YAP/TAZ-Dependent Reprogramming of Colonic Epithelium Links ECM Remodeling to Tissue Regeneration. Cell Stem Cell 22, 35-49.e7.

Zheng, Y., Valdez, P.A., Danilenko, D.M., Hu, Y., Sa, S.M., Gong, Q., Abbas, A.R., Modrusan, Z., Ghilardi, N., Sauvage, F.J. De, et al. (2008). Interleukin-22 mediates early host defense against attaching and effacing bacterial pathogens. 14, 282–289.

Zhou, A.-X., Hartwig, J.H., and Akyürek, L.M. (2010). Filamins in cell signaling, transcription and organ development. Trends Cell Biol. 20, 113–123.

Zwiggelaar et al., 2020 (2020). LSD1 represses a neonatal/reparative gene program in adult intestinal epithelium. BioRxiv 2020.02.20.958363.

